# First mitogenome phylogeny of the sun bear *Helarctos malayanus* reveals a deep split between Indochinese and Sundaic lineages

**DOI:** 10.1101/2022.09.14.507900

**Authors:** Miriam N. Kunde, Axel Barlow, Achim Klittich, Aliya Yakupova, Riddhi P. Patel, Jörns Fickel, Daniel W. Förster

## Abstract

The sun bear *Helarctos malayanus* is one of the most endangered ursids, and to date classification of sun bear populations has been based almost exclusively on geographic distribution and morphology. The very few molecular studies focusing on this species were limited in geographic scope. Using archival and non-invasively collected sample material, we have added a substantial number of complete or near-complete mitochondrial genome sequences from sun bears of several range countries of the species’ distribution. We here report 32 new mitogenome sequences representing sun bears from Cambodia, Thailand, Peninsular Malaysia, Sumatra and Borneo. Reconstruction of phylogenetic relationships revealed two matrilines that diverged ∼290 thousand years ago: one restricted to portions of mainland Indochina (China, Cambodia, Thailand; “Mainland clade”), and one comprising bears from Borneo, Sumatra, Peninsular Malaysia but also Thailand (“Sunda clade”). Generally recent coalescence times in the mitochondrial phylogeny suggest that recent or historical demographic processes have resulted in a loss of mtDNA variation. Additionally, analysis of our data in conjunction with shorter mtDNA sequences revealed that the Bornean sun bear, classified as a distinct subspecies (*H. m. euryspilus*), does not harbour a distinctive matriline. Further molecular studies of *H. malayanus* are needed, which should ideally include data from nuclear loci.

## INTRODUCTION

Archival and non-invasively collected sample material is increasingly utilized in molecular studies of wildlife species, particularly if these are rare, elusive, protected, or inhabit areas that are difficult to access (e.g. Paijmans et al., 2020; Mengüllüoglu et al. 2021; Hessels et al. 2021; Sacks et al. 2021; von Thaden et al. 2021; Cho et al. 2022). DNA extracted from such material is usually highly degraded, rendering downstream analyses difficult (e.g. Pääbo 1989). However, targeted capture coupled with high throughput sequencing enables retrieval of genetic information from such degraded sample material, with mitochondrial DNA (mtDNA) often the marker of choice, because the high copy number of mitochondria per cell benefits sequence recovery (e.g. Paijmans et al. 2013; Jones & Good, 2016). MtDNA sequences enriched by targeted capture can then be used to resolve population genetic structure and taxonomic relationships within species, and can provide insights into the historical processes that have shaped contemporary populations (Avise, 2000).

With regard to threatened species, conservation management greatly benefits from such knowledge (Wilting et al. 2011; Wilting et al. 2015; Martins et al. 2017a), as observational data or phenotypic characters may not accurately reflect how genetic variation is partitioned within species (Patel et al. 2017; Wilting et al. 2016; Martins et al. 2017b), or which populations have evolved independently for long periods of time (‘evolutionary significant units’, ESUs). Such information is still lacking for many wildlife species, including those inhabiting ecosystems in global biodiversity hotspots, such as Southeast (SE) Asia, that are increasingly adversely affected by anthropogenic land-use and land-cover change (Tilker et al 2020, Nguyen et al 2021).

Such a knowledge gap also exists for the Malayan sun bear *Helarctos malayanus*. In 1999, the sun bear was identified as the “least studied bear species” and has earned the moniker ‘the forgotten’ or ‘neglected’ bear (Meijaard & Nooteboom 1999). Research neglect poses a threat to the conservation of sun bears and it has been argued that they should be given research priority within the bear family *Ursidae* (Servheen 1999). However, this call for action has been ignored for almost two decades, and sun bears are still too understudied for effective conservation efforts to be implemented (Kunde, 2017; Kunde et al 2020). These small bears were once widely distributed across SE Asia (Fig. 1), occurring in eastern India, Bangladesh, Myanmar, Thailand, Laos, Cambodia, Vietnam, Malaysia, Indonesia, and southern China (Fitzgerald and Krausman 2002). Over the past decades, their distribution area has dramatically shrunk, and sun bears have lost 30-60% of their habitat over the past 30 years (Scotson et al. 2017), with a concomitantly severe decline in population size. They now occur mostly in small patches of primary lowland and rainforest remnants (Scotson et al. 2017). As a result of habitat loss and population size decline, the species has become listed as *Vulnerable* on the IUCN Red List of threatened species (Scotson et al. 2017) and is in Appendix 1 of CITES (CITES, 2022).

**Figure 1:**
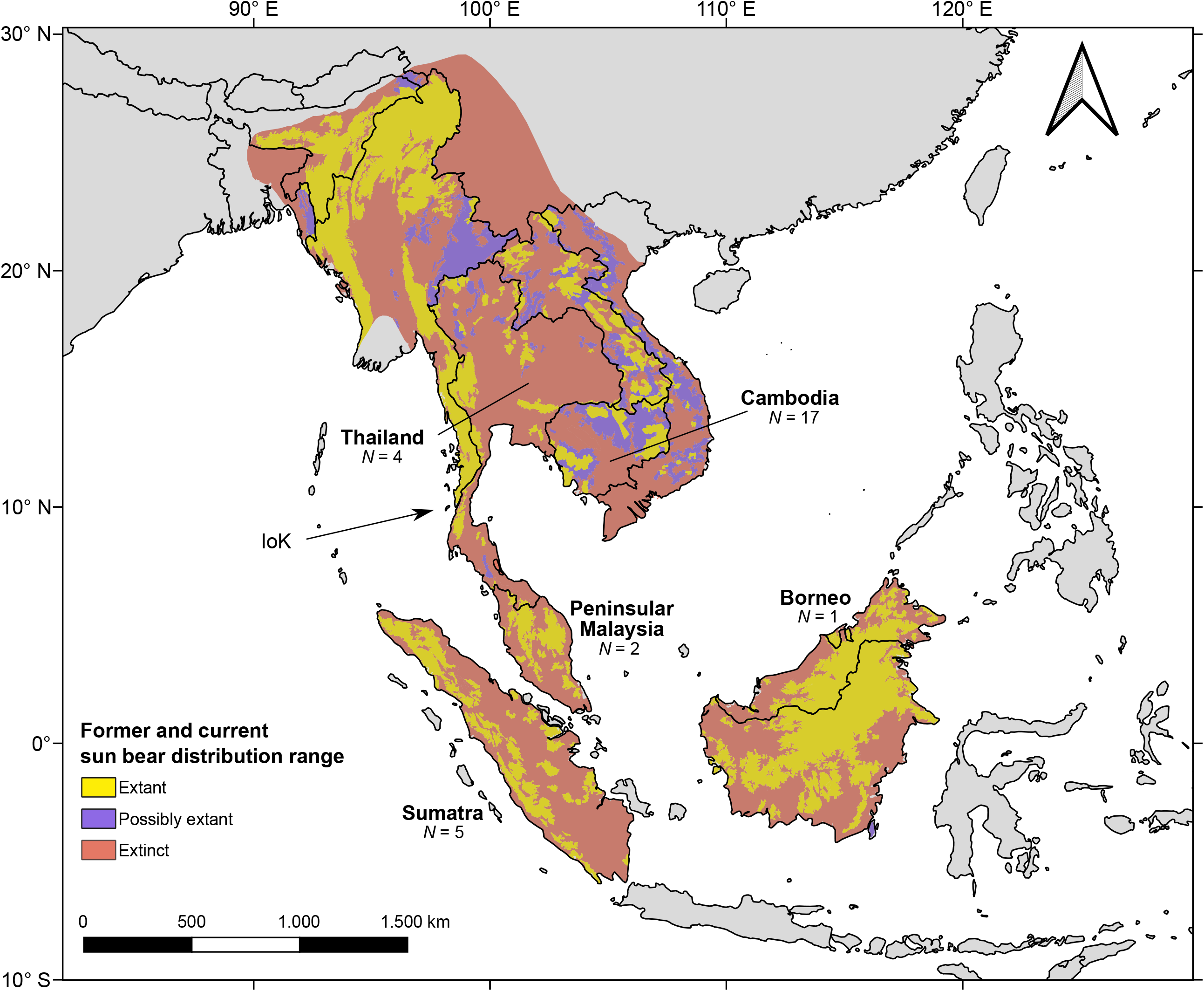
Former and current *H. malayanus* distribution range (following Scotson et al. 2017). The number of samples for which we were able to retrieve mitochondrial genomes is indicated, as is the Isthmus of Kra (“IoK”).

Sun bears are threatened by many factors such as excessive hunting, habitat loss and fragmentation, but also by inconsistent wildlife law enforcement policy in range countries (Meijaard & Nooteboom 1999; Fredriksson 2005; INTERPOL 2014; Krishnasamy & Shepherd 2014, Scotson et al. 2017, Kunde et al. 2020). Despite being strictly protected globally since 1979 (Servheen 1999; Wong et al. 2004; Augeri 2005; Shepherd & Shepherd 2010), sun bears are still considered game species in some countries (Servheen,1999, Loke et al. 2020). In addition, cubs are hunted for the illegal pet trade (Foley et al. 2011; Krishnasamy & Shepherd 2014; Lee et al. 2015; Gomez et al. 2020), and body parts are used for making wine, soup, lucky charms, and traditional Chinese medicine products (Scotson & Hunt 2008; Scotson & Downie 2009; Shepherd & Shepherd 2010; Cantlay et al. 2017). The massive decline of population size, deficiency in life history data, lack of knowledge about the species’ current distribution, high mortality rates, and diminishing population health strongly suggest that current conservation strategies for sun bears are failing to achieve desired results (Schneider et al. 2020).

Molecular studies are needed to describe and manage the remaining genetic diversity in sun bears. A phylogeographic approach can reveal management and evolutionary units (Avise 1992; Moritz 1994), help to identify conservation priorities (Goossens et al. 2013), and pinpointing suitable reintroduction locations (Apollonio et al. 2014).

To date, studies of *H. malayanus* using molecular markers have been limited in geographic scope (Onuma et al. 2006; Kunde et al. 2020; Lai et al 2021) and we thus lack knowledge about the distribution of intraspecific genetic diversity across this species’ range. To address this shortcoming, we sequenced mitochondrial genomes from samples across a large portion of the sun bear distribution, utilizing both archival material from natural history museums as well as non-invasively collected material (saliva). Using these complete mitochondrial genomes (mitogenomes), we elucidated the geographic distribution of maternal lineages (matrilines) and provide insights into the species’ history.

## MATERIALs AND METHODS

### Samples

Dried tissue remains from the skull or tissue from the nasal cavity of 29 archival sun bear specimens were collected from several natural history museums (Supplementary Table S1), providing a broad geographic distribution of samples.

We also collected saliva samples from 21 sun bears, which originated from 12 provinces in Cambodia and which were housed at the ‘Free the Bears’ sanctuary in Phnom Tamao Zoo and Wildlife Rescue Centre. We included up to two samples per province in order to obtain a balanced geographic coverage of Cambodia.

### Archival and non-invasively collected DNA

DNA from archival samples was extracted following Wilting et al. (2016), in a laboratory designed for and limited to the use of archival material. DNA extractions were conducted in batches consisting of three samples and one negative control and verified to be of sun bear origin by amplification of a 146 bp long portion of the mitochondrial cytochrome *b* gene using species specific primers (Supplemental Table S2).

DNA extractions from non-invasively collected samples (salivary mucosal cells) were carried out using the GEN-IAL First DNA *All*-tissue DNA extraction Kit (GEN-IAL GmbH, Troisdorf, Germany) following manufacturer’s instructions. This work was carried out in a separate facility, dedicated to molecular work with non-invasively collected material.

### Library preparation and hybridization capture

Illumina sequencing libraries were constructed following Fortes & Paijmans (2015), using double-indexing with 8-bp indexes. Prior to indexing we conducted qPCRs to determine the optimal PCR cycle number for each sample.

To reduce effects by target DNA degradation and external DNA contaminations, we carried out hybridisation capture for both sample types (archival, non-invasively collected).

Baits for hybridization capture were generated using long range (LR) PCR products that spanned three large overlapping regions of the mitogenome (using a fresh *Helarctos malayanus* sample as template). The LR-PCR primers were designed using the genbank reference sequence NC_009968: HMA1-F (5’
s-ACGACCTCGATGTTGGATCAGG-3’) and HMA1-R (5’- AGGGCTACAGCGAACTCGAGA-3’), yielding a 6152 bp long product, HMA2-F (5’- GCCACACTCATTCACACCTACCA-3’) and HMA2-R (5’-AGTCCTTTCTGGTTGGAGACTGTG- 3’), yielding a 5299 bp long product, and HMA3-F (5’-ACCAACGCCTGAGCCCTACT-3’) and HMA3-R (5’-GCGCTTTAGTGAGGGAGGCC-3’), yielding a 6416 bp long product. Amplifications were carried out in 50µL reaction volumes: 18µL dH_2_O, 25µL Bioline Myfi mix (Bioline GmbH, Germany), 2µL of forward and reverse primers (10μM), 3µL of template DNA (100ng/μL). PCR conditions were as follows: initial denaturation at 94°C for 3 min, followed by 40 cycles of denaturation at 94°C for 30 s; annealing at 60°C for 30 s; extension at 68°C for 7 min with the final extension at 68° C for 10 min.

LR-PCR products were sheared (target size 300bp), pooled equimolarly and then converted into biotinylated baits following Maricic et al. (2010). Targeted capture was carried out using hybridization temperatures appropriate for sample type (Paijmans et al. 2016). Paired-end sequencing was carried out on the Illumina MiSeq platform (Illumina, San Diego, CA, USA) using v3 150-cycle kits.

### Bioinformatic workflow

We de-multiplexed paired-end reads using BCL2FASTQ v2.17.1.14 (Illumina, Inc.) and removed adapter sequences using CUTADAPT v1.3 (Martin 2011). Using TRIMMOMATIC (Bolger, Lohse, and Usadel 2014), we applied a sliding window approach for quality trimming with the phred quality threshold set at Q=20. Next, we merged the adapter-clipped and quality-trimmed sequences using FLASH v1.2.8 (Magoc and Salzberg 2011). Merged sequences were then mapped to the reference sun bear mitogenome sequence (GenBank accession no. NC_009968) with the BWA aln algorithm v0.7.10 (Li and Durbin 2009). Mapped sequences were then de-duplicated using MARKDUPLICATES from PICARD-TOOLS v1.106 (https://github.com/broadinstitute/picard), followed by variant calling using SAMTOOLS v1.1 (Li et al. 2009) and BCFTOOLS v1.2 (http://github.com/samtools/bcftools). Positions along the mitogenome were N-masked if their sequence depth was below 5×, or when they did not conform to an 80% majority rule for base calling.

### Summary statistics

The final data set of complete or near-complete sun bear mitogenomes consisted of 15 archival and 17 non-invasively collected samples, analyzed together with two published sequences (GenBank: FM177765 and EF196664; see Table 1 for details).

**Table 1.** Sample details

| Sample ID | Region | Sample type | raw reads | clipped,<br>trimmed<br>and merged<br>sequences | deduplicated<br>sequences<br>mapped<br>to<br>mitogenome | %<br>mitogenome<br>covered $\geq 5x$ |
| --- | --- | --- | --- | --- | --- | --- |
| HMA_1_MA | Peninsular Malaysia | archival | 962,556 | 395,594 | 7,632 | 97.95% |
| HMA_2_MA | Peninsular Malaysia | archival | 4,279,758 | 1,958,381 | 325,844 | 99.92% |
| HMA_4_TH | Thailand | archival | 782,922 | 357,378 | 7,550 | 95.96% |
| HMA_5_SU | Sumatra | archival | 655,610 | 300,916 | 74,221 | 98.26% |
| HMA_13 | *unknown* | archival | 703,722 | 319,130 | 43,562 | 96.89% |
| HMA_15_TH | Thailand | archival | 1,999,180 | 881,484 | 172,478 | 98.64% |
| HMA_16_SU | Sumatra | archival | 1,431,992 | 643,106 | 56,190 | 98.49% |
| HMA_17_SU | Sumatra | archival | 1,214,110 | 548,477 | 20,344 | 97.89% |
| HMA_19_BO | Borneo | archival | 998,120 | 407,017 | 28,629 | 98.87% |
| HMA_24_TH | Thailand | archival | 1,105,638 | 499,533 | 7,984 | 97.72% |
| HMA_26_TH | Thailand | archival | 1,119,718 | 521,872 | 149,064 | 98.71% |
| HMA_021_CAM | Cambodia | saliva | 632,824 | 285,401 | 72,500 | 98.96% |
| HMA_027_CAM | Cambodia | saliva | 842,154 | 349,635 | 34,873 | 99.03% |
| HMA_028_CAM | Cambodia | saliva | 853,854 | 366,541 | 10,990 | 98.77% |
| HMA_031_CAM | Cambodia | saliva | 707,374 | 273,830 | 8,936 | 98.74% |
| HMA_035_CAM | Cambodia | saliva | 1,311,174 | 578,748 | 16,942 | 98.71% |
| HMA_037_CAM | Cambodia | saliva | 1,230,298 | 547,022 | 7,365 | 98.12% |
| HMA_042_CAM | Cambodia | saliva | 860,156 | 362,656 | 12,187 | 98.81% |
| HMA_046_CAM | Cambodia | saliva | 1,816,556 | 766,221 | 36,603 | 98.89% |
| HMA_052_CAM | Cambodia | saliva | 1,623,318 | 585,438 | 10,331 | 98.71% |
| HMA_057_CAM | Cambodia | saliva | 992,996 | 441,147 | 8,488 | 98.73% |
| HMA_061_CAM | Cambodia | saliva | 1,619,226 | 608,738 | 46,909 | 99.02% |
| HMA_065_CAM | Cambodia | saliva | 2,232,478 | 822,778 | 28,069 | 99.09% |
| HMA_069_CAM | Cambodia | saliva | 723,998 | 321,510 | 23,069 | 98.87% |
| HMA_079_CAM | Cambodia | saliva | 951,314 | 419,079 | 58,623 | 99.03% |
| HMA_086_CAM | Cambodia | saliva | 961,508 | 421,987 | 7,186 | 95.14% |
| HMA_087_CAM | Cambodia | saliva | 954,048 | 375,369 | 17,300 | 98.99% |
| HMA_095_CAM | Cambodia | saliva | 1,073,632 | 370,083 | 8,575 | 98.80% |
| HMA_34002_SU | Sumatra | archival | 2,380,950 | 1,083,797 | 139,932 | 98.84% |
| HMA_34004_SU | Sumatra | archival | 5,117,344 | 1,934,439 | 58,458 | 99.32% |
| HMA_15638 | *unknown* | archival | 2,897,942 | 936,442 | 25,587 | 98.27% |
| HMA_A5351 | *unknown* | archival | 4,472,384 | 1,667,180 | 19,935 | 98.49% |
| FM177765 | *unknown* | *Genbank* | n.a. | n.a. | n.a. | n.a. |
| EF196664 | China | *Genbank* | n.a. | n.a. | n.a. | n.a. |

Further sample details are provided in Suppl. Table S1.

Mitogenome sequences were aligned and manually curated in Geneious v. 8.1.9 (Kearse et al. 2012). Summary statistics for the complete dataset (*N* = 34), as well as subdivisions based on phylogenetic analyses (below) were generated using DnaSP v.6.12.03 (Rozas et al. 2017). For these estimates we excluded sites with missing data, ambiguous data, or gaps.

### Phylogenetic analyses

To generate a time-calibrated mitochondrial phylogeny of sun bears, we first estimated the root age in a species level analysis calibrated based on the fossil ages of other representatives of the Ursidae. We then applied this root age estimate to a population level analysis of all sun bear mitochondrial sequences. This two step strategy was chosen to best accommodate the assumptions of the tree priors used in these respective analyses.

For the species level analysis, we aligned two divergent sun bear mitogenomes (HMA_26_TH, and HMA_35_CAMB) and one mitogenome from each of the following nine species: *Ursus maritimus* (Polar bear; acc. no. AF303111), *Ursus arctos* (Brown bear; acc. no. HQ685956), *Ursus thibetanus* (Asian black bear; acc. no. EF196661), *Ursus americanus* (American black bear; acc. no. AF303109), *Melursus ursinus* (Sloth bear; acc. no. FM177763), *Tremarctos ornatus* (Spectacled bear; acc. no. EF196665), *Ailuropoda melanoleuca* (Giant Panda; acc. no. EF196663), *Arctodus simus* (extinct Short-faced bear; acc. no. FM177762) and *Ursus ingressus* (extinct Cave bear; acc. no. KX641331). We then used PartitionFinder v1.1.1 (Lanfear et al. 2012) to identify an optimal set of partitions and substitution models among all combinations of tRNAs, rRNAs and the three codon positions of the protein coding genes, using the greedy search algorithm and considering all substitution models available in BEAST, using the Bayesian Information Criterion (BIC). The Bayesian phylogenetic analysis package BEAST v.1.8.2 (Drummond et al. 2012) was then used to estimate species phylogeny and divergence times. A birth-death tree model was used, with a lognormal relaxed clock model, with a uniform prior on the mean substitution rate of 0 to 20 percent per million years (My). Time-calibration was achieved using informative uniform priors based on fossil and other evidence, following the approach of a previous study of the Ursidae (Kumar et al. 2017). Monophyly was enforced for all calibrated nodes. These were: (1) a prior based on the divergence of the Tremarctinae and Ursinae clades between 7 to 14 My, based on a fossil of Tremarctine bear *Plionarctos* (Tedford, 2009); (2) a prior for the basal divergence of the Ursidae with an upper limit of 12 My, based on a fossil of the Ailuropodinae (Abella et al. 2012) and a lower limit of 20 My, determined by a molecular dating study (Wu et al. 2015); (3) a prior on divergence of the Ursinae between 4.3 – 6 My based on the age of *Ursus minimus* (Gustafson, 1978); and (4) a prior on the divergence of *U. arctos* and *U. maritimus* between 0.48 My – 1.1 My based on previous studies (Hailer et al. 2012; Li et al. 2011; Cahill et al. 2015). All other priors were left at default values. Initial runs showed a lack of convergence for some substitution model parameters, and so we substituted them with simpler models (Supplementary Table S3). The final BEAST run involved 20 million generations, sampling the MCMC chain every 1,000 generations, in order to achieve adequate burn in and posterior sampling of all parameters (ESS > 200), assessed using the program Tracer 1.7.

The divergence estimate recovered for the two sun bear mitogenomes (mean 0.304 My, standard deviation 0.0376 My) was then applied as normal prior on the root height for the intraspecific analyses, and were run with a Bayesian Skyline coalescent population model with 8 groups. Data partitions and substitutions were estimated using PartitionFinder as described previously. As we did not expect variation in mitochondrial substitution rate within sun bears, we used a strict clock model with a uniform prior on the per-lineage substitution rate of 0 to 20 percent per My. Details of the MCMC run were described previously. The maximum clade credibility tree was then selected from the posterior sample, and node heights scaled to the median posterior age, using TreeAnnotator 1.8.2. A Bayesian skyline plot was generated using Tracer.

We also generated a dataset combining portions of our mitochondrial sequences with a set of recently published ∼1,800bp long mtDNA sequences from sun bears from Thailand, Peninsular Malaysia and Borneo (GenBank acc. no. MW316324 - MW316405; Lai et al. 2021). This combined dataset of shorter sequences consisted of 116 mtDNA sequences: 32 from this study, 82 from Lai et al. (2021), one from Yu et al. (2007), and one from Krause et al. (2008) (Suppl. Table S4). The alignment had a length of 1,466bp after removal of the repetitive portion of the control region. To visualize the relationship among these sequences, we generated a median joining (MJ) network using POPART v.1.7. (Bandelt et al.1999).

## RESULTS

Using targeted capture coupled with high throughput sequencing, we were able to recover near complete mitogenome sequences with ≥5× sequencing depth at each base for 32 samples, 15 from archival and 17 from non-invasively collected samples (Table 1; Genbank accession numbers: XXX-YYY). Without considering alignment gaps and ambiguous base calls (i.e. Ns in sequences), we detected 25 unique haplotypes among the 32 mitogenomes. Putative haplotypes were mostly shared among sun bears from Cambodia (Suppl. Table S5), however, one such putative haplotype was shared by bears from Thailand and Borneo (HMA_15_TH, HMA_19_BOR). Summary statistics for the complete dataset (*N =* 34), as well as subdivisions of the data based on phylogenetic analyses (below) are presented in Table 2.

**Table 2.**
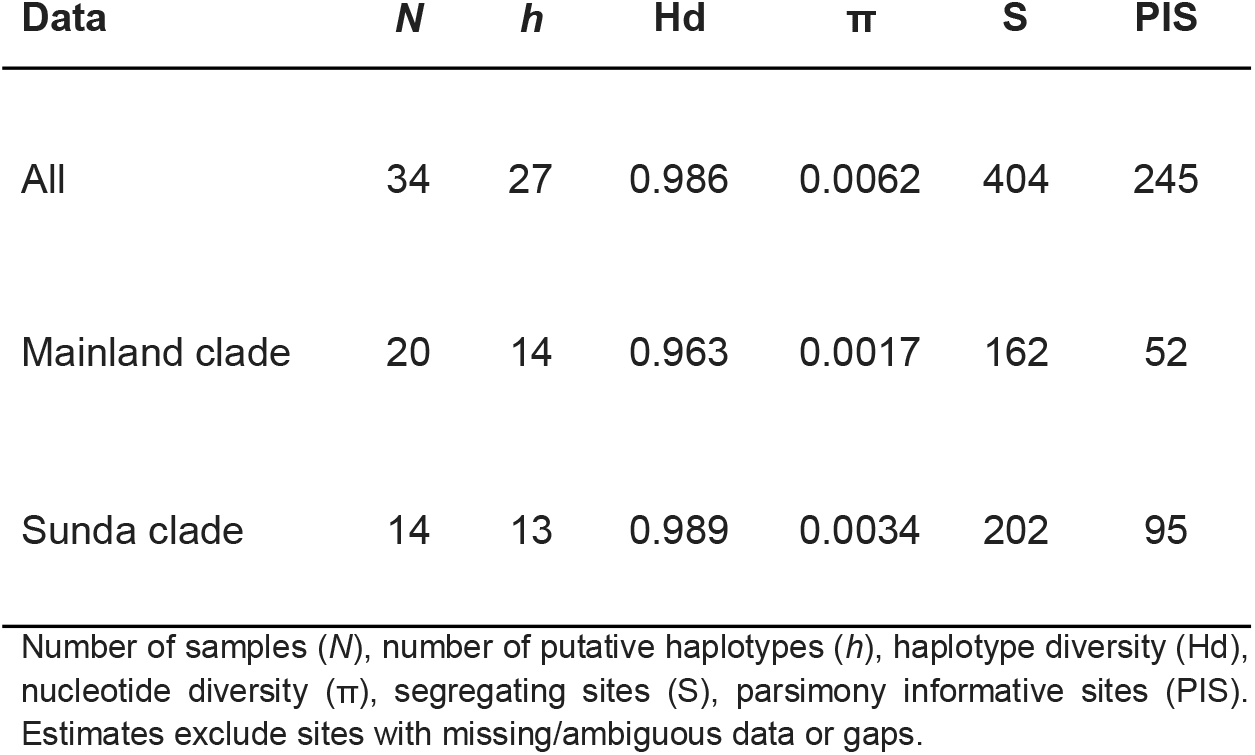
Summary statistics for sun bear mitogenome sequences

| Data | *N* | *h* | Hd | $\pi$ | S | PIS |
| --- | --- | --- | --- | --- | --- | --- |
| All | 34 | 27 | 0.986 | 0.0062 | 404 | 245 |
| Mainland clade | 20 | 14 | 0.963 | 0.0017 | 162 | 52 |
| Sunda clade | 14 | 13 | 0.989 | 0.0034 | 202 | 95 |

Number of samples (*N*), number of putative haplotypes (*h*), haplotype diversity (Hd), nucleotide diversity ( $\pi$ ), segregating sites (S), parsimony informative sites (PIS). Estimates exclude sites with missing/ambiguous data or gaps.

### Mitogenome Phylogeny

Phylogenetic analysis of the sun bear mitogenome dataset (34 sequences) revealed two well-supported clades (posterior clade credibility 1 for each clade), henceforth referred to as “Mainland clade” and “Sunda clade”, that diverged ∼290 thousand years ago (kya; CI_95_: 365-215 kya) (Fig. 2).

**Figure 2:**
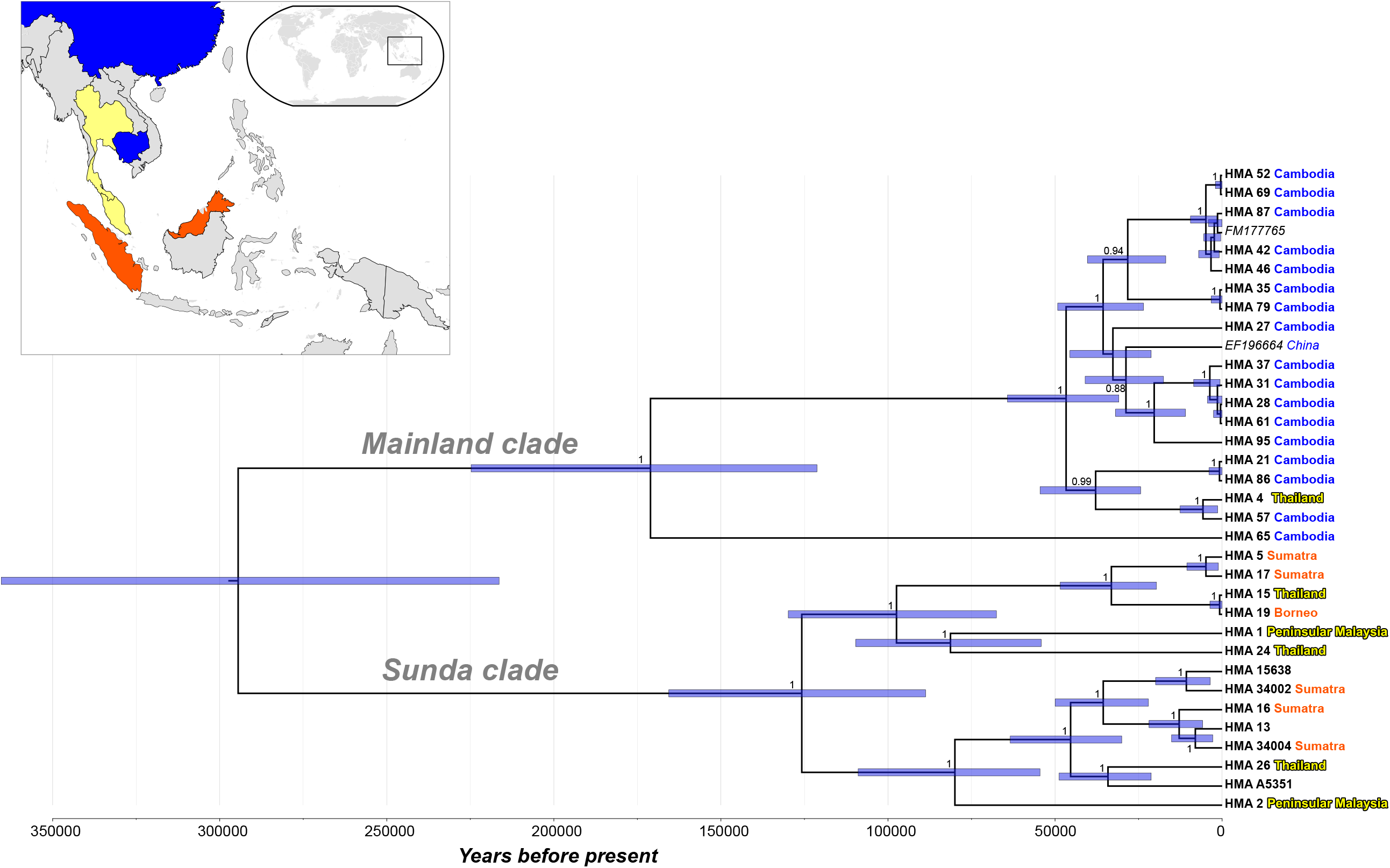
Bayesian phylogenetic tree reconstructed using *H. malayanus* mitogenome sequences (*N* = 34). Two main branches are labeled according to clade assignment. Sequences are labeled with sample ID and geographic origin (if known); text colour of geographic origin follows the inset map (top left). Blue bars indicate 95% credibility intervals for coalescence times in years.

The Mainland clade includes sequences (*N* = 20) from Cambodia, China, and Thailand, as well as a sequence from a sun bear of unknown provenance (Genbank acc. no. FM177765; Krause et al. 2008). Within this clade, two lineages diverged ∼170 kya (CI_95_: 225-120 kya), of which one is currently only represented by a single sequence from Cambodia. The remaining 19 sequences are in the second lineage, with a coalescence-time estimated at ∼45 kya (CI_95_: 60-30 kya). These sequences are mostly derived from Cambodian samples (16 of 19), and no obvious geographic structuring is apparent.

The Sunda clade includes sequences (*N* = 14) from Borneo, Sumatra and Peninsular Malaysia (all part of Sundaland), but also from Thailand, and from three archival samples of unknown provenance (sample details in Suppl Table S1). This clade also gave rise to two major lineages, which diverged ∼125 kya (CI_95_: 165-85 kya), and whose samples have a large geographic overlap: both include sun bears from Sumatra, Peninsular Malaysia and Thailand. Notably, we were only able to retrieve the mitogenome for one sample from Borneo, so we are unable to determine if both or only one of these lineages occurs on this island. Furthermore, as mentioned above, the sample from Borneo shared the same putative haplotype with a sample from Thailand.

It is evident that both mitogenome clades are present in Thailand. Unfortunately, it is not known whether these Thai sun bear samples originated from North or South of the Isthmus of Kra (5-13° N), an important zoogeographic transition zone between Sundaland and continental mainland.

### Shorter mtDNA sequences

We also considered the 34 mitogenome sequences in the context of a recent phylogenetic study on shorter mtDNA sequences of sun bears from Thailand (it is unknown if these samples originated from North or South of the Isthmus of Kra), Peninsular Malaysia and Borneo (Lai et al. 2021) and constructed a median-joining (MJ) network from this combined dataset (*N =* 116 sequences; alignment length 1,466 bp; Figure 3).

**Figure 3:**
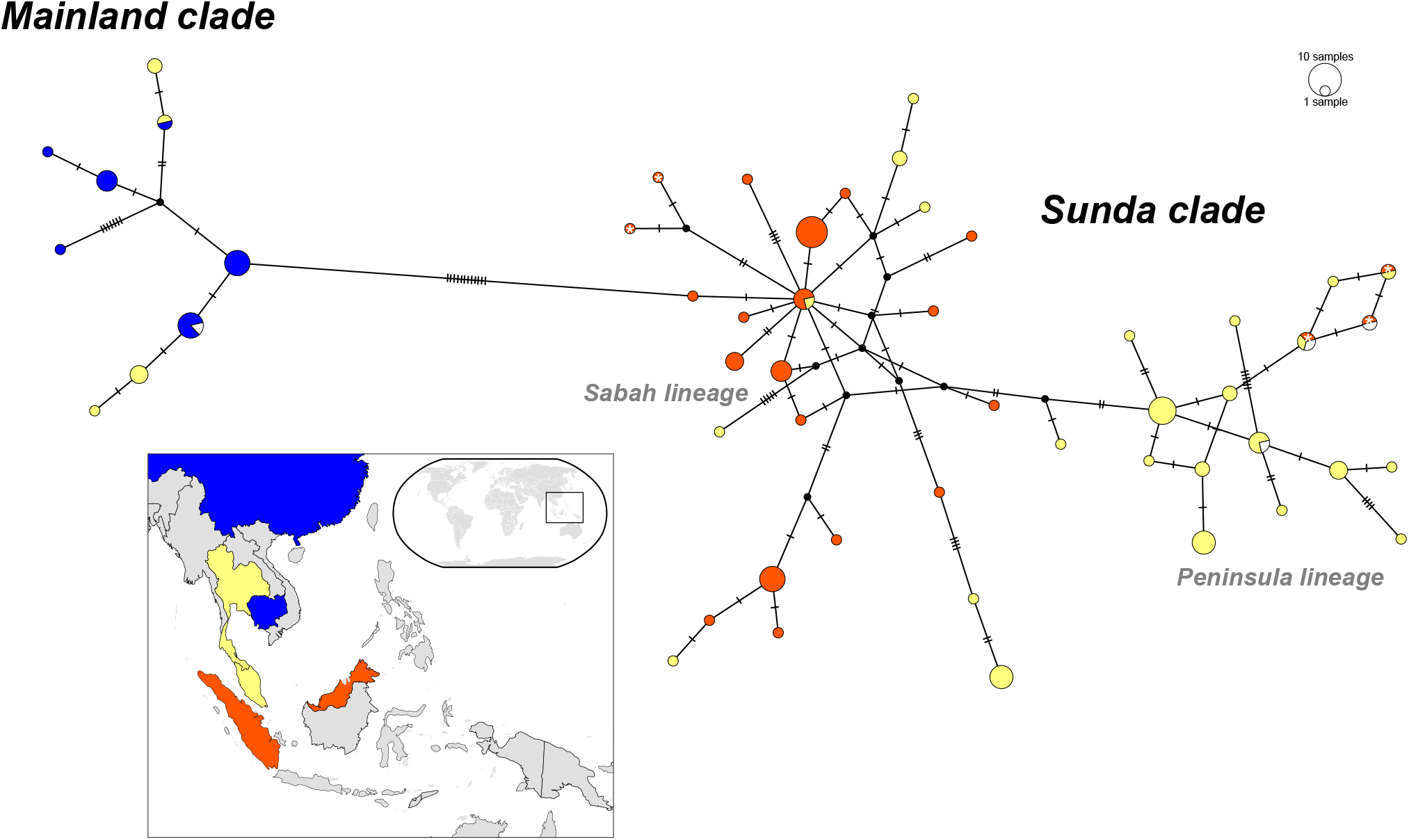
Median-joining network constructed using shorter mtDNA sequences (*N =* 116; 1,466 bp). The number of mutations separating connected haplotypes is indicated by the number of dashes along the connecting line. Black labels indicate the two major mtDNA clades; grey labels indicate lineages previously reported by Lai et al. (2021). Colouring of haplotypes is according to geographic origin (if known), following inset map (bottom left). The five haplotypes from Sumatra are indicated by white asterisks (‘*’).

Consistent with the mitogenome phylogeny, both clades, Mainland and Sunda, are apparent in the MJ network, whereby the Mainland clade was enlarged by Thailand samples T7, T8, T9, T10 and T14 (from Lai et al. 2021).

Genetic structuring is apparent in the Sunda clade, which is consistent with previous findings (Lai et al. 2021). Samples originating mostly from Peninsular Malaysia formed a lineage (“Peninsula Lineage” in Lai et al., 2021; Fig. 3) different from the one formed by samples of the Sundaic islands (“Sabah lineage” (all from Borneo) in Lai et al., 2021; Fig. 3). However, this structuring is not geographically exclusive. For example, our samples from Sumatra (HMA_16_SU, HMA_34002_SU, HMA_34004_SU) and Thailand (HMA_26_TH) are part of the Peninsula lineage, together with samples from Thailand (T2, T6, T11, T16, and T17; Lai et al. 2021). Our samples from Peninsular Malaysia (HMA_1_MA, HMA_2_MA), Borneo (HMA_19_BO) and Sumatra (HMA_5_SU, HMA_17_SU) are included in the Sabah lineage, together with a sample from Thailand (HMA_24_TH). This lineage was already reported to include sequences from Thailand (T1, T3, T4, T5, T13, T18) and Peninsular Malaysia (P15, P17, P27) by Lai et al. (2021). Thus, both lineages are found in Peninsular Malaysia, Thailand and Sumatra, while the Sabah lineage is also present on Borneo.

The Peninsula and Sabah lineages correspond to the two lineages within the Sunda clade we identified using the complete mitogenomes (Fig. 2).

## DISCUSSION

By incorporating archival and non-invasively collected material in our sampling, and using targeted capture coupled with high throughput sequencing, we successfully retrieved data for mitochondrial genomes of sun bears from a large portion of their geographic distribution.

We identified two matrilines within the sampled distribution of the species: one restricted to portions of mainland SE Asia (encompassing China, Cambodia and Thailand), while the other occurs in Sundaland (Peninsular Malaysia, Sumatra and Borneo), as well as parts of the mainland (Thailand). Despite our efforts, the failure of some archival samples to yield results means that our coverage of the sun bear distribution in mainland SE Asia is unfortunately limited; we are missing data for Vietnam, Laos, Myanmar, and India.

It has been suggested that sun bears have originated in what is now Peninsular Malaysia and Borneo, and that the divergence of observed sun bear morphological forms commenced in the Pliocene (Meijaard 2004). The coalescence time for the two clades indicates that sun bear populations diverged during the Middle Pleistocene, a much younger date than the proposed Pliocene divergence. Fossil evidence places sun bears in Indochina during the Middle Pleistocene (Louys, 2012), while evidence for the species’ presence on the Sundaic islands is dated to the late Pleistocene (Medway 1964; Long, et al. 1996; Tougard 2001), although an earlier presence is conceivable as the mammalian fossil record for Sumatra and Borneo is sparse prior to the Late Pleistocene (Meijaard 2004). While molecular estimates of divergence times need to be considered with caution, our mitogenome data do suggest the presence of sun bears on the Sundaic islands by the late Middle Pleistocene.

It could be argued that the coalescence time for Sundaic sun bears of ∼125 kya is not consistent with the origin of sun bears in this region (i.e. Peninsular Malaysia and Borneo, following Meijaard, 2004), but this estimate only takes into account mitochondrial variation of extant populations in that region. It should also be noted that without the retrieval of the divergent mitogenome lineage within the Mainland clade from sample HMA_65_CAM (Fig. 2), the coalescence time estimate for sun bears of the Mainland clade would have been only ∼45 kya, much younger than the Sundaic expansion. Further sampling throughout the sun bear distribution may improve our understanding of the species’ origin, as additional mitogenome lineages may be unsampled thus far (e.g. Mengüllüoğlu et al. 2021). However, while the reconstructed mitochondrial phylogeny in general is very recent, historical and recent demographic processes may have resulted in the loss of variation (mitochondrial lineages) that existed in the past, thereby obscuring older evolutionary processes. Thus, the question of the sun bears’ origin is likely best addressed using nuclear markers. The issue is further compounded by possible secondary contact of sun bears in the region where the species may have originated.

### Secondary contact

The geographic distribution of mitogenome clades reflects vicariance between sun bears of mainland Indochina and Sundaland, a biogeographic pattern that is observed in many other SE Asian species that are widely distributed (Woodruff & Turner 2009). It is unknown if the Mainland clade was lost in sun bears before, during, or after colonization of the Sundaland. If this loss occurred before or during colonization, this suggests that sun bears from the Sunda clade have persisted in the Thai-Malay Peninsula since the Middle Pleistocene. If the loss of the Mainland clade occurred after colonization, this suggests secondary contact of the two lineages in the Thai-Malay Peninsula, as both major clades are found in Thailand. Such migration was likely possible due to the exposed shelf that connected the mainland and the major islands of the Sundaland during the low sea levels of the late Pleistocene (Bird et al. 2005), and has been inferred for other large mammals, such as leopard *Panthera pardus* (Wilting et al. 2016), leopard cat *Prionailurus bengalensis* (Patel et al. 2017), and muntjac *Muntiacus muntjak* (Martins et al. 2017b).

If there is secondary contact of these two lineages following a northward migration of sun bears of the Sunda clade, we can consider two scenarios: (1) the replacement of sun bears of the Mainland clade by bears of the Sunda clade in portions of the Thai-Malay Peninsula, or (2) the recolonization of the Thai-Malay Peninsula by bears from the Sunda clade following a local extinction of Mainland clade sun bears, possibly resulting from the Toba volcano super eruption on Sumatra ∼74 kya (Costa et al., 2014). The first scenario suggests that Sundaic sun bears would have had some form of competitive advantage over individuals of the local mainland population during the period(s) when the Sunda Shelf was exposed, and migration of Sundaic bears to the Thai-Malay Peninsula was possible. In the second scenario, we would need to consider that the eruption would have most likely also severely affected the Sumatran sun bear population (as suggested for other species, e.g. leopard: Wilting et al. 2016). In this context, it is of note that the Peninsula lineage (following Lai et al., 2021) has thus far only been detected in Sumatra, Peninsular Malaysia and Thailand, but not on Borneo, raising the question where this lineage may have endured following the Toba eruption. A candidate region is Kalimantan, the Indonesian portion of Borneo, for which no mtDNA data is currently available.

Clearly, the currently available molecular data is not adequate to address these open questions, and further sampling is needed. Analyses should ideally also include nuclear genomic data, which would also be extremely valuable to address the taxonomic uncertainty regarding the sun bears of Borneo.

### Bornean sun bears

There are clear phenotypic differences between the sun bears on Borneo and those from elsewhere in the species’ range, and currently two subspecies are recognized: the broadly distributed *H. m. malayanus* and the Bornean *H. m. euryspilus* (Meijaard, 2004).

The rainforest on Borneo is less fruit-bearing than the rainforest on Sumatra or on the Asian mainland, and this reduced food availability may have favored smaller body size and modified dentition and skull morphology on Borneo (Meijaard 2004; Wich et al. 2011). However, it has been argued that these differences reflect a phenotypic response to food availability and foraging behavior, potentially misleading taxonomy, as a well-nourished bear may simply grow bigger than a malnourished bear (Kitchener 2010).

Mitochondrial data can not disclose the mechanisms behind the observed phenotypic differences. However, it does show that Bornean sun bears harbor one Sundaic lineage (Sabah lineage) that is present throughout the region, including Sumatra, Peninsular Malaysia and Thailand, while a second widespread Sundaic lineage (Peninsula lineage) has thus far not been observed on Borneo. If the absence of the Peninsula lineage on Borneo is true (and not the result of sampling bias) then it is unclear why the two lineages do not coexist on Borneo. It is also unknown how far their divergence time (∼125 kya) relates to the divergence of Bornean sun bears. In this context, it can also not be discounted that historical or modern human-mediated translocation of sun bears may have contributed to the observed distribution of mitochondrial lineages in the Sundaic region, further complicating matters.

### Human-mediated translocations

Sun bears are hunted for the illegal pet trade (Foley et al. 2011; Krishnasamy & Shepherd 2014; Lee et al. 2015; Gomez et al. 2020), and are traded in and among the Sundaic islands (e.g. Gomez et al. 2019). Rescue and release of such trafficked bears may result in the introduction of non-endemic mtDNA haplotypes to local populations. It is worth noting that such translocations do not need to be recent. There is a history of human-mediated introduction or translocation of mammals (e.g. Long et al. 2003) for various reasons (e.g. cultural, commercial, pest-control, accidental). Such translocations included large mammals and carnivores in SE Asia, among others cervids (*Rusa timorensis* and *R. unicolor*, Martins et al. 2017a), pigs (Groves, 1984), the leopard cat (*Prionailurus bengalensis*, Patel et al. 2017), the Malay civet (*Viverra tangalunga*, Veron et al 2014), and the Asian palm civet (*Paradoxurus hermaphroditus*, Flannery et al. 1995). Because some of these human-mediated translocations had commercial reasons (e.g. Groves, 1984), it is conceivable that there was also historical trade in bears and their derivatives (e.g. Hose, 1893), which may have included the translocation and subsequent (intentional or unintentional) release of sun bears. Investigation of Early Holocene samples using ancient DNA techniques would be valuable in addressing the impact of human-mediated translocations. Unfortunately, sun bear fossils are rare, and the species inhabits an environment that is not conducive to ancient DNA preservation (e.g. Paijmans et al. 2013).

### Concluding remarks

Thus far, the classification of sun bear populations has been based on geography and morphological traits, the latter having only been investigated using a small dataset. It is important to corroborate/supplement such efforts using molecular studies (Kitchener 2010). Our study reveals a clear and deep split into two matrilines that divide sun bears from the Sundaic region (Borneo, Sumatra, Peninsular Malaysia, Thailand) from others on the Indochinese mainland (China, Cambodia, Thailand). As has been observed in mtDNA studies of other SE Asian vertebrates, these Sundaic and mainland lineages are both present in the Thai-Malay Peninsula. With regards to sun bear taxonomy, we observe that the Bornean subspecies *H. m. euryspilus* does not carry a distinctive mitochondrial lineage, but rather one that is present throughout Sundaland (Borneo, Sumatra, Peninsular Malaysia, Thailand). To improve our understanding of sun bear evolution and biogeography, further molecular studies are needed. Two obvious priorities are to include samples from a border geographical context and to obtain data from nuclear genomes. Until such research has been carried out, we encourage ex-situ conservation management to take the existence of the two highly differentiated matrilines into account.

## Supporting information

Supplementary Material for this publication

## Data Accessibility Statement

We have deposited the primary data underlying these analyses as follows:

- Mitochondrial genome sequences: GenBank acc. no. XXX-YYY
- Sampling locations, haplotypes, and collection details are in Supplementary Table S1
- Cytochrome B primer sequences to verify target species are in Supplementary Table S2
- Sequences used in analysis of short mtDNA sequence are in Supplementary Table S4

## Competing Interests Statement

None.

## Author Contributions

M.N.K. and D.W.F. conceived of the study. M.N.K. and A.W. collected samples and obtained permits. M.N.K. and R.P.P. conducted laboratory procedures. M.N.K., A.B., A.K., A.Y., R.P.P. and D.W.F. conducted analyses. M.N.K. and D.W.F. wrote the paper with input from all authors.

## Acknowledgements

We would like to thank the Kingdom of Cambodia and the Australian government for issuing permits (CITES export permit No KH0936, CITES PWS2014-AU-002163 and PWS2015-AU-000922, multiple use import permit PWS2014-AU-001113), and Griffith University for ethics approval for this research (ethics permits GU Ref No: ENV/04/13/AEC, and GU Ref No: ENV/02/13/AEC). We would also like to thank Free the Bears Cambodia, State Museum of Natural History Stuttgart (SMNS), Zoologisches Forschungsmuseum Alexander Koenig Bonn (ZMFK), Museum of Natural History Vienna (NHM), Museum für Tierkunde Dresden (MTD), National Museum of Natural History Paris (MNHN), Naturmuseum Senckenberg Frankfurt (SNG), Bavarian State Collection of Zoology Munich (ZSM), and Museum für Naturkunde Berlin (ZMB) for allowing sample collection, and Andreas Wilting for sampling the archival samples. MK thanks Anke Schmidt for laboratory supervision and support, and Tim Heupink for assisting in sample transport. This work was funded by the Leibniz-Association grant SAW-2013-IZW-2.

## Notes

### Competing Interest Statement

The authors have declared no competing interest.

