## Supplementary Material for this publication for "First mitogenome phylogeny of the sun bear *Helarctos malayanus* reveals a deep split between Indochinese and Sundaic lineages"

| Contents |  |
| --- | --- |
| Suppl. Table S1: detailed sample info | Pg. 1-4 |
| Suppl. Table S2: cytochrome b primers & PCR details | Pg. 4 |
| Suppl. Table S3: Substitution models (BEAST) | Pg. 5 |
| Suppl. Table S4: sample details for analysis of short mtDNA sequences | Pg. 6-8 |
| Suppl. Table S5: shared mitogenome haplotypes | Pg. 8 |

**Suppl. Table S1**: detailed sample info

| **Sample ID** | **Collection Location** | **Museum or  Sanctuary  Location** | **Mueum/**  **Sanctuary** | **Museum or Sanctuary ID** | **collection year** | **Provided by** | **Sample type** |
| --- | --- | --- | --- | --- | --- | --- | --- |
| HMA_1_MA | Jahore, Malaysia | Stuttgart, Germany | SMNS | 21754 | 1897 | Dr. Stefan Merker, Staatliches Museum für Naturkunde Stuttgart | nasal bone |
| HMA_2_MA | Jahore, Malaysia | Stuttgart, Germany | SMNS | 21753 | 1897 | Dr. Stefan Merker, Staatliches Museum für Naturkunde Stuttgart | nasal bone |
| HMA_3_IN | Burma/Vorderasien | Bonn, Germany | ZFMK | ZFMK 32.21 | *unknown* | Dr. Jan Decher, Zoologisches Forschungsinstitut und Museum Alexander Koenig, Bonn | skin |
| HMA_4_TH | Siam (Thailand) | Bonn, Germany | ZFMK | ZFMK 81.55 | 1921 | Dr. Jan Decher, Zoologisches Forschungsinstitut und Museum Alexander Koenig, Bonn | tissue |
| HMA_5_SU | prov. Deli, Nord-Sumatra | Bonn, Germany | ZFMK | ZFMK 65.5 | 1963/64 | Dr. Jan Decher, Zoologisches Forschungsinstitut und Museum Alexander Koenig, Bonn | bone (vertebra) |
| HMA_6_JA | Java, Indonesia | Wien, Austria | NHM | B 4249 | Oct, 1929 | Dr. habil. Frank Emmanuel Zachos, Naturhistorisches Museum Wien | Skin (fur) |
| HMA_7_SU | Sumatra | Dresden, Germany | MTD | B6076 | 1827 | Dr. Clara Stefen, Senckenberg Naturhistorische Sammlungen Dresden | Skin+ hair |
| HMA_8_VI | Vietnam | Paris, France | MNHN | 1929-430 | *unknown* | Geraldine Veron, Museum National d'Histoire Naturelle, Paris | tissue from skull |
| HMA_9_JA | Java, Indonesia | Paris, France | MNHN | A-2132 | *unknown* | Geraldine Veron, Museum National d'Histoire Naturelle, Paris | tissue from skull |
| HMA_10_CH | Ho-Chi-Minh-city/Saigon | Frankfurt, Germany | SNG | 5790 | *unknown* | Katrin Krohmann, Senckenberg Naturkundemuseum, Frankfurt | skin |
| HMA_11_SU | Sunda Islands, SE-Asia | Frankfurt, Germany | SNG | 15461 | *unknown* | Katrin Krohmann, Senckenberg Naturkundemuseum, Frankfurt | skin |
| HMA_12_SU | Sunda Islands, SE-Asia | Frankfurt, Germany | SNG | 15776 | *unknown* | Katrin Krohmann, Senckenberg Naturkundemuseum, Frankfurt | skin |
| HMA_13 | Zoo animal | Frankfurt, Germany | SNG | 15777 | *unknown* | Katrin Krohmann, Senckenberg Naturkundemuseum, Frankfurt | dry tissue from skin |
| HMA_14_SU | Lampong, South-Sumatra | Munich, Germany | ZSM | 1908/539 | *unknown* | Michael Hiermeier, Zoologische Staatssammlung München | loose bones (tissue) |
| HMA_15_TH | Siam (Thailand) | Munich, Germany | ZSM | 1918/31 | *unknown* | Michael Hiermeier, Zoologische Staatssammlung München | Skeleton (tissue) |
| HMA_16_SU | Medan Deli, east coast Sumatra | Munich, Germany | ZSM | 1918/482 | *unknown* | Michael Hiermeier, Zoologische Staatssammlung München | Skull (nasal bone) |
| HMA_17_SU | Medan Deli, east coast Sumatra | Munich, Germany | ZSM | 1910/154 | *unknown* | Michael Hiermeier, Zoologische Staatssammlung München | Skull (nasal bone) |
| HMA_18_BO | Borneo | Munich, Germany | ZSM | 1907/231 | *unknown* | Michael Hiermeier, Zoologische Staatssammlung München | Skull (nasal bone) |
| HMA_19_BO | Borneo | Munich, Germany | ZSM | 1907/485 | *unknown* | Michael Hiermeier, Zoologische Staatssammlung München | Skull (nasal bone) |
| HMA_20_BO | Borneo | Munich, Germany | ZSM | 1907/471 | *unknown* | Michael Hiermeier, Zoologische Staatssammlung München | Skull (nasal bone) |
| HMA_21_BO | Borneo | Munich, Germany | ZSM | 1907/638 | *unknown* | Michael Hiermeier, Zoologische Staatssammlung München | Skull (nasal bone) |
| HMA_22_TH | Siam (Thailand) | Munich, Germany | ZSM | 1904/169 | May-04 | Michael Hiermeier, Zoologische Staatssammlung München | Skull (nasal bone) |
| HMA_23_TH | Siam (Thailand) | Munich, Germany | ZSM | 1906/143 | *unknown* | Michael Hiermeier, Zoologische Staatssammlung München | Skull (nasal bone) |
| HMA_24_TH | Siam (Thailand) | Munich, Germany | ZSM | 1906/142 | *unknown* | Michael Hiermeier, Zoologische Staatssammlung München | Skull (nasal bone) |
| HMA_25_TH | Siam (Thailand) | Munich, Germany | ZSM | 1911/19 | 09/03/1911 | Michael Hiermeier, Zoologische Staatssammlung München | Skull (nasal bone) |
| HMA_26_TH | Siam (Thailand) | Munich, Germany | ZSM | 1906/149 | *unknown* | Michael Hiermeier, Zoologische Staatssammlung München | Skull (nasal bone) |
| HMA_013_CAM | Siem Reap (Cambodia) | Pnom Penh, Cambodia | FtB | Ranee | 2014 | Free the Bears/ Kingdom of Cambodia | Saliva |
| HMA_028_CAM | Kompong Cham (Cambodia) | Pnom Penh, Cambodia | FtB | Mien | 2014 | Free the Bears/ Kingdom of Cambodia | Saliva |
| HMA_035_CAM | Siem Reap (Cambodia) | Pnom Penh, Cambodia | FtB | Tonle Sap | 2014 | Free the Bears/ Kingdom of Cambodia | Saliva |
| HMA_042_CAM | Koh Kong (Cambodia) | Pnom Penh, Cambodia | FtB | Kong | 2014 | Free the Bears/ Kingdom of Cambodia | Saliva |
| HMA_057_CAM | Koh Kong (Cambodia) | Pnom Penh, Cambodia | FtB | Ooshie | 2014 | Free the Bears/ Kingdom of Cambodia | Saliva |
| HMA_079_CAM | Oddar Meanchey (Cambodia) | Pnom Penh, Cambodia | FtB | Sam | 2014 | Free the Bears/ Kingdom of Cambodia | Saliva |
| HMA_086_CAM | Prey Vihear (Cambodia) | Pnom Penh, Cambodia | FtB | Win | 2014 | Free the Bears/ Kingdom of Cambodia | Saliva |
| HMA_031_CAM | Oddar Meanchey (Cambodia) | Pnom Penh, Cambodia | FtB | Phat | 2014 | Free the Bears/ Kingdom of Cambodia | Saliva |
| HMA_052_CAM | Kompong Speu (Cambodia) | Pnom Penh, Cambodia | FtB | Hasa | 2014 | Free the Bears/ Kingdom of Cambodia | Saliva |
| HMA_061_CAM | Preah Vihear (Cambodia) | Pnom Penh, Cambodia | FtB | Hope-bee | 2014 | Free the Bears/ Kingdom of Cambodia | Saliva |
| HMA_065_CAM | Preah Vihear (Cambodia) | Pnom Penh, Cambodia | FtB | Jacko | 2014 | Free the Bears/ Kingdom of Cambodia | Saliva |
| HMA_080_CAM | Kratie (Cambodia) | Pnom Penh, Cambodia | FtB | Kratie | 2014 | Free the Bears/ Kingdom of Cambodia | Saliva |
| HMA_090_CAM | Stung Treng (Cambodia) | Pnom Penh, Cambodia | FtB | Sybil Sunbeam | 2014 | Free the Bears/ Kingdom of Cambodia | Saliva |
| HMA_021_CAM | Kompong Thom (Cambodia) | Pnom Penh, Cambodia | FtB | Ju-ju | 2014 | Free the Bears/ Kingdom of Cambodia | Saliva |
| HMA_027_CAM | Kratie (Cambodia) | Pnom Penh, Cambodia | FtB | Dodo | 2014 | Free the Bears/ Kingdom of Cambodia | Saliva |
| HMA_037_CAM | Ratanakiri (Cambodia) | Pnom Penh, Cambodia | FtB | Buddy | 2014 | Free the Bears/ Kingdom of Cambodia | Saliva |
| HMA_046_CAM | Kompong Thom (Cambodia) | Pnom Penh, Cambodia | FtB | Kiem | 2014 | Free the Bears/ Kingdom of Cambodia | Saliva |
| HMA_087_CAM | Kompong Som (Cambodia) | Pnom Penh, Cambodia | FtB | Deena | 2014 | Free the Bears/ Kingdom of Cambodia | Saliva |
| HMA_088_CAM | Ratanakiri (Cambodia) | Pnom Penh, Cambodia | FtB | Abbie | 2014 | Free the Bears/ Kingdom of Cambodia | Saliva |
| HMA_095_CAM | Pursat (Cambodia) | Pnom Penh, Cambodia | FtB | Holly | 2014 | Free the Bears/ Kingdom of Cambodia | Saliva |
| HMA_069_CAM | Stung Treng (Cambodia) | Pnom Penh, Cambodia | FtB | Po Sary | 2014 | Free the Bears/ Kingdom of Cambodia | Saliva |
| HMA_34002_SU | Sumatra | Berlin, Germany | ZMB | ZMB_MA_34002 | *unknown* | Dr. Frieder Mayer, Naturkundemuseum Berlin | Skull |
| HMA_34004_SU | Sumatra | Berlin, Germany | ZMB | ZMB_MA_34004 | *unknown* | Dr. Frieder Mayer, Naturkundemuseum Berlin | Skull |
| HMA_17531_TH | Thailand | Berlin, Germany | ZMB | ZMB_MA_17531 | *unknown* | Dr. Frieder Mayer, Naturkundemuseum Berlin | Skull |
| HMA_15638 | *unknown* | Berlin, Germany | ZMB | ZMB_MA_15638 | *unknown* | Dr. Frieder Mayer, Naturkundemuseum Berlin | Skull |
| HMA_A5351 | *unknown* | Berlin, Germany | ZMB | ZMB_MA_A5351 | *unknown* | Dr. Frieder Mayer, Naturkundemuseum Berlin | Skull |
| HMA_17245_BO | Borneo | Berlin, Germany | ZMB | ZMB_MA_17245 | *unknown* | Dr. Frieder Mayer, Naturkundemuseum Berlin | Skull |

**Suppl. Table S2**: cytochrome b primers & PCR details

| **Primers** | F: ACACCGAAATCTTTCTCACT  R: AAGGAAATAAAATGCTCGGAGAC |
| --- | --- |
| **PCR mix** | Reaction volume of 15 µl:  6.48 µl of dH2O, 3 µl GoTaq Buffer, 1.2 µl MgCl2 (25 mM), 1 µl BSA (20mM), 0.25 µl dNTPs (10mM), 0.5 µl Primer F (10mM), 0.5 µl Primer R (10mM), 0.075 µl GoTaq Polym (5 U/ µl), 2 µl DNA template |
| **Thermocycler conditions** | Initial denaturation at  95° C for 2 min, then 40 cycles of denaturion at 95° C for 30 s, annealing at 59° C for 30 s, and extension at 72° C for 30 s, with a final extension at 72° C for 5 min. |

**Suppl. Table S3:** Substitution models (BEAST)

| **SPECIES LEVEL ANALYSIS** | | |  |
| --- | --- | --- | --- |
| ***Partition*** | ***BIC Model*** | ***Used model*** | ***mtDNA regions*** |
| 1 | TrN+G | TrN+G | 12s, 16s_p1, 16s_pt2, Ala, Arg, Asn, Asp, Cys, Gln, Glu, Gly, His, Ile, Leu2, Lys, ND6_CP2, Ser2, Thr, Trp, Tyr |
| 2 | TrN+I+G | TrN+G | ATP8_ATP6_CP3, COX2_CP1, CYTB_CP1, Leu1, ND1_CP1, ND2_CP1, ND3_CP1, ND4L_CP1, ND4_CP1, ND5_CP1, Pro, Ser1, Val |
| 3 | HKY+I+G | HKY+I+G | COX1_CP2, COX2_CP2, COX3_CP2, CYTB_CP2, ND1_CP2, ND2_CP2, ND3_CP2, ND4L_CP2, ND4_CP2 |
| 4 | TrN+I+G | TrN+I+G | COX3_CP3, CYTB_CP3, ND1_CP3, ND2_CP3, ND3_CP3, ND4L_CP3, ND4_CP3, ND5_CP3, ND6_CP3 |
| 5 | K80+I | K80+I | COX1_CP1, COX3_CP1, Met |
| 6 | TrN+G | TrN+G | ATP8_ATP6_CP2, COX1_CP3, COX2_CP3 |
| 7 | TrN+I+G | TrN+G | ATP8_ATP6_CP1, ND5_CP2, ND6_CP1 |
| 8 | HKY+G | HKY+G | D-loop, Phe |
| **POPULATION LEVEL ANALYSIS** | | |  |
| ***Partition*** | ***BIC Model*** | ***Used model*** | ***mtDNA regions*** |
| 1 | K80 | K80 | 16s_pt2, COX1_CP1, COX2_CP1, COX3_CP1, CYTB_CP1, Cys, Met, ND1_CP1, ND3_CP1, ND4L_CP1, Ser1, Trp |
| 2 | HKY+G | HKY+G | ATP8_ATP6_CP2, COX1_CP3, COX2_CP3, D-loop, Gly, Leu1, Pro |
| 3 | HKY | HKY | ATP8_ATP6_CP1, COX1_CP2, COX2_CP2, COX3_CP2, CYTB_CP2, ND1_CP2, ND2_CP2, ND3_CP2, ND4L_CP2, ND4_CP2, ND5_CP2, Tyr |
| 4 | TrN | TrN | COX3_CP3, CYTB_CP3, ND1_CP3, ND2_CP3, ND3_CP3, ND4L_CP3, ND4_CP3, ND5_CP3, ND6_CP1, ND6_CP3 |
| 5 | TrN | TrN | 12s, 16s_p1, ATP8_ATP6_CP3, Ala, Arg, Asn, Asp, Gln, Glu, His, Ile, Leu2, Lys, ND2_CP1, ND4_CP1, ND5_CP1, ND6_CP2, Phe, Ser2, Thr, Val |

**Suppl. Table S4:** sample details for analysis of short mtDNA sequences

| **Sample ID** | **Region** | **Sample Type** | **Accession Number** | **Study** |
| --- | --- | --- | --- | --- |
| HMA_1_MA | Peninsular Malaysia | archival | X | *this study* |
| HMA_2_MA | Peninsular Malaysia | archival | X | *this study* |
| HMA_4_TH | Thailand | archival | X | *this study* |
| HMA_5_SU | Sumatra | archival | X | *this study* |
| HMA_13 | *unknown* | archival | X | *this study* |
| HMA_15_TH | Thailand | archival | X | *this study* |
| HMA_16_SU | Sumatra | archival | X | *this study* |
| HMA_17_SU | Sumatra | archival | X | *this study* |
| HMA_19_BO | Borneo | archival | X | *this study* |
| HMA_24_TH | Thailand | archival | X | *this study* |
| HMA_26_TH | Thailand | archival | X | *this study* |
| HMA_021_CAM | Cambodia | saliva | X | *this study* |
| HMA_027_CAM | Cambodia | saliva | X | *this study* |
| HMA_028_CAM | Cambodia | saliva | X | *this study* |
| HMA_031_CAM | Cambodia | saliva | X | *this study* |
| HMA_035_CAM | Cambodia | saliva | X | *this study* |
| HMA_037_CAM | Cambodia | saliva | X | *this study* |
| HMA_042_CAM | Cambodia | saliva | X | *this study* |
| HMA_046_CAM | Cambodia | saliva | X | *this study* |
| HMA_052_CAM | Cambodia | saliva | X | *this study* |
| HMA_057_CAM | Cambodia | saliva | X | *this study* |
| HMA_061_CAM | Cambodia | saliva | X | *this study* |
| HMA_065_CAM | Cambodia | saliva | X | *this study* |
| HMA_069_CAM | Cambodia | saliva | X | *this study* |
| HMA_079_CAM | Cambodia | saliva | X | *this study* |
| HMA_086_CAM | Cambodia | saliva | X | *this study* |
| HMA_087_CAM | Cambodia | saliva | X | *this study* |
| HMA_095_CAM | Cambodia | saliva | X | *this study* |
| HMA_34002_SU | Sumatra | archival | X | *this study* |
| HMA_34004_SU | Sumatra | archival | X | *this study* |
| HMA_15638 | *unknown* | archival | X | *this study* |
| HMA_A5351 | *unknown* | archival | X | *this study* |
| FM177765 | *unknown* | *-* | FM177765 | Krause et al. 2008 |
| EF196664 | China | *-* | EF196664 | Yu et al. 2007 |
| P1 | Peninsula/Johor | Blood on FTA card | MW316360 | Lai et al. 2021 |
| P2 | Peninsula/Pahang | Blood on FTA card | MW316361 | Lai et al. 2021 |
| P3 | Peninsula/Pahang | Blood on FTA card | MW316362 | Lai et al. 2021 |
| P4 | Peninsula/Perak | Blood on FTA card | MW316363 | Lai et al. 2021 |
| P5 | Peninsula/Perak | Blood on FTA card | MW316364 | Lai et al. 2021 |
| P6 | Peninsula/Selangor | Blood on FTA card | MW316365 | Lai et al. 2021 |
| P7 | Peninsula/- | Blood on FTA card | MW316366 | Lai et al. 2021 |
| P8 | Peninsula/Pahang | Blood on FTA card | MW316367 | Lai et al. 2021 |
| P9 | Peninsula/Pahang | Blood on FTA card | MW316368 | Lai et al. 2021 |
| P10 | Peninsula/Pahang | Incisor | MW316369 | Lai et al. 2021 |
| P11 | Peninsula/Penang | Hair | MW316370 | Lai et al. 2021 |
| P12 | Peninsula/Pahang | Hair | MW316371 | Lai et al. 2021 |
| P13 | Peninsula/Johor | Hair | MW316372 | Lai et al. 2021 |
| P14 | Peninsula/Perak | Hair | MW316373 | Lai et al. 2021 |
| P15 | Confiscated | Hair | MW316374 | Lai et al. 2021 |
| P16 | Peninsula/Kedah | Hair | MW316375 | Lai et al. 2021 |
| P17 | Confiscated | Hair | MW316376 | Lai et al. 2021 |
| P18 | Peninsula/- | Muscle | MW316377 | Lai et al. 2021 |
| P19 | Peninsula/Terengganu | Muscle | MW316378 | Lai et al. 2021 |
| P20 | Peninsula/Zoo | Hair | MW316379 | Lai et al. 2021 |
| P21 | Peninsula/Zoo | Hair | MW316380 | Lai et al. 2021 |
| P22 | Peninsula/Terengganu | Hair | MW316381 | Lai et al. 2021 |
| P23 | Peninsula/Zoo | Hair | MW316382 | Lai et al. 2021 |
| P24 | Peninsula/Pahang | Hair | MW316383 | Lai et al. 2021 |
| P25 | Peninsula/Selangor | Hair | MW316384 | Lai et al. 2021 |
| P26 | Peninsula/Melaka | Blood in EDTA | MW316385 | Lai et al. 2021 |
| P27 | Peninsula/Zoo | Blood in EDTA | MW316386 | Lai et al. 2021 |
| P28 | Peninsula/Zoo | Blood in EDTA | MW316387 | Lai et al. 2021 |
| B1 | Sabah/Keningau | Hair | MW316324 | Lai et al. 2021 |
| B2 | Sabah/— | Hair | MW316325 | Lai et al. 2021 |
| B3 | Sabah/Lahad Datu | Hair | MW316326 | Lai et al. 2021 |
| B4 | Sabah/Pitas | Hair | MW316327 | Lai et al. 2021 |
| B5 | Sabah/— | Hair | MW316328 | Lai et al. 2021 |
| B6 | Sabah/Tawau | Hair | MW316329 | Lai et al. 2021 |
| B7 | Sabah/Kota Kinabalu | Hair | MW316330 | Lai et al. 2021 |
| B8 | Sabah/Tawau | Hair | MW316331 | Lai et al. 2021 |
| B9 | Sabah/Sipitang | Hair | MW316332 | Lai et al. 2021 |
| B10 | Sabah/Penampang | Hair | MW316333 | Lai et al. 2021 |
| B11 | Sabah/Penampang | Hair | MW316334 | Lai et al. 2021 |
| B12 | Sabah/Ranau | Hair | MW316335 | Lai et al. 2021 |
| B13 | Sabah/Kuamut | Hair | MW316336 | Lai et al. 2021 |
| B14 | Sabah/Kudat | Hair | MW316337 | Lai et al. 2021 |
| B15 | Sabah/Kinabatangan | Hair | MW316338 | Lai et al. 2021 |
| B16 | Sabah/Ranau | Hair | MW316339 | Lai et al. 2021 |
| B17 | Sabah/— | Hair | MW316340 | Lai et al. 2021 |
| B18 | Sabah/— | Hair | MW316341 | Lai et al. 2021 |
| B19 | Sabah/Kota Marudu | Hair | MW316342 | Lai et al. 2021 |
| B20 | Sabah/Sipitang | Hair | MW316343 | Lai et al. 2021 |
| B21 | Sabah/Tawau | Hair | MW316344 | Lai et al. 2021 |
| B22 | Sabah/— | Hair | MW316345 | Lai et al. 2021 |
| B23 | Sabah/Kudat | Hair | MW316346 | Lai et al. 2021 |
| B24 | Sabah/Sipitang | Hair | MW316347 | Lai et al. 2021 |
| B25 | Sabah/Tawau | Hair | MW316348 | Lai et al. 2021 |
| B26 | Sabah/Tawau | Hair | MW316349 | Lai et al. 2021 |
| B27 | Sabah/Keningau | Hair | MW316350 | Lai et al. 2021 |
| B28 | Sabah/Labuk dan Sugut | Hair | MW316351 | Lai et al. 2021 |
| B29 | Sabah/— | Hair | MW316352 | Lai et al. 2021 |
| B30 | Sabah/Nabawan | Hair | MW316353 | Lai et al. 2021 |
| B31 | Sabah/— | Hair | MW316354 | Lai et al. 2021 |
| B32 | Sabah/Kota Marudu | Hair | MW316355 | Lai et al. 2021 |
| B33 | Sabah/Kudat | Hair | MW316356 | Lai et al. 2021 |
| B34 | Sabah/Labuk dan Sugut | Hair | MW316357 | Lai et al. 2021 |
| B35 | Sabah/Labuk dan Sugut | Hair | MW316358 | Lai et al. 2021 |
| B36 | Sabah/Tawau | Hair | MW316359 | Lai et al. 2021 |
| T1 | Thailand/West | Hair | MW316388 | Lai et al. 2021 |
| T2 | Thailand/— | Hair | MW316389 | Lai et al. 2021 |
| T3 | Thailand/— | Hair | MW316390 | Lai et al. 2021 |
| T4 | Thailand/— | Hair | MW316391 | Lai et al. 2021 |
| T5 | Thailand/— | Hair | MW316392 | Lai et al. 2021 |
| T6 | Thailand/— | Hair | MW316393 | Lai et al. 2021 |
| T7 | Thailand/East | Hair | MW316394 | Lai et al. 2021 |
| T8 | Thailand/— | Hair | MW316395 | Lai et al. 2021 |
| T9 | Thailand/— | Hair | MW316396 | Lai et al. 2021 |
| T10 | Thailand/— | Hair | MW316397 | Lai et al. 2021 |
| T11 | Thailand/West | Hair | MW316398 | Lai et al. 2021 |
| T12 | Thailand/East | Hair | MW316399 | Lai et al. 2021 |
| T13 | Thailand/— | Hair | MW316400 | Lai et al. 2021 |
| T14 | Thailand/— | Hair | MW316401 | Lai et al. 2021 |
| T15 | Thailand/— | Hair | MW316402 | Lai et al. 2021 |
| T16 | Thailand/— | Hair | MW316403 | Lai et al. 2021 |
| T17 | Thailand/— | Hair | MW316404 | Lai et al. 2021 |
| T18 | Thailand/— | Hair | MW316405 | Lai et al. 2021 |

**Suppl. Table S5**: shared mitogenome haplotypes

HMA_52_CAMB, HMA_69_CAMB

HMA_87_CAMB, HMA_46_CAMB

HMA_35_CAMB, HMA_79_CAMB

HMA_28_CAMB, HMA_31_CAMB, HMA_61_CAMB

HMA_86_CAMB, HMA_21_CAMB

HMA_15_TH, HMA_19_BO
